# A Metagenomic Meta-Analysis Reveals Functional Signatures of Health and Disease in the Human Gut Microbiome

**DOI:** 10.1101/286419

**Authors:** Courtney Armour, Stephen Nayfach, Katherine Pollard, Thomas Sharpton

**Affiliations:** Molecular and Cellular Biology Program, Oregon State University, Corvallis, OR, USA; Department of Microbiology, Oregon State University, Corvallis, OR, USA; Department of Energy Joint Genome Institute, Walnut Creek, California, USA; Gladstone Institutes, San Francisco, CA, USA; Department of Epidemiology & Biostatistics, Institute for Human Genetics, Quantitative Biology Institute, and Institute for Computational Health Sciences, University of California, San Francisco, CA, USA; Chan-Zuckerberg Biohub, San Francisco, CA, USA; Department of Statistics, Oregon State University, Corvallis, OR, USA

## Abstract

While recent research indicates that human health depends, in part, upon the symbiotic relationship between gut microbes and their host, the specific interactions between host and microbe that define health are poorly resolved. Metagenomic clinical studies clarify this definition by revealing gut microbial taxa and functions that stratify healthy and diseased individuals. However, the typical single-disease focus of microbiome studies limits insight into which microbiome features robustly associate with health, indicate general deviations from health, or predict specific diseases. Additionally, the focus on taxonomy may limit our understanding of how the microbiome relates to health given observations that different taxonomic members can fulfill similar functional roles. To improve our understanding of the association between the gut microbiome and health, we integrated about 2,000 gut metagenomes obtained from eight clinical studies in a statistical meta-analysis. We identify characteristics of the gut microbiome that associate generally with disease, including functional alpha-diversity, beta-diversity, and beta-dispersion. Moreover, we resolve microbiome modules that stratify diseased individuals from controls in a manner independent of study-specific effects. Many of the differentially abundant functions overlap multiple diseases suggesting a role in host health, while others are specific to a single disease and may associate with disease-specific etiologies. Our results clarify potential microbiome-mediated mechanisms of disease and reveal features of the microbiome that may be useful for the development of microbiome-based diagnostics. Ultimately, our study clarifies the definition of a healthy microbiome and how perturbations to it associate with disease.

## INTRODUCTION

Mounting evidence implicates the gut microbiome as a critical component of human health. For example, research demonstrates that the gut microbiome contributes to immunity, nutrition, and behavior[1, 2]. Accordingly, gut microbiomes of diseased individuals tend to harbor different taxa and encode different genes than those of healthy individuals[3]. These observations motivate the hypothesis that human health depends, in part, upon the taxonomic composition and biological functions executed by the gut microbiome. Therefore, researchers have sought to identify the properties of the human gut microbiome that signify health and disease. Such signatures are valuable to resolve because they provide important context for the development of disease diagnostics, clarify disease etiology, and generate insight into how microbiomes should be amended to restore health.

An extensive array of prior investigations focused on defining the characteristics of the gut microbiome that signify health or disease. One such effort was led by the Human Microbiome Project, which defined the structure and function of the gut microbiome that associates with clinically healthy, urban, North Americans[4]. Other investigations used clinical 16S data to determine how the structure of the gut microbiome of diseased individuals differs from that of healthy individuals. More recently, a smaller set of investigations used shotgun metagenomes to resolve how both the structure and functional diversity of the gut microbiome associates with disease[5–12]. However, almost all of these investigations have focused on a single disease population and a matching control. Very few studies integrate data across multiple populations, incorporate data from other studies, or compare patterns across various disease types. Consequently, it is unclear which associations are robust to population or study effects. Moreover, we possess limited insight into which associations are specific to a disease-type versus those that are common to myriad diseases. These limitations hinder our ability to develop robust clinical diagnostics from microbiome data and obscure our understanding of the potential mechanisms through which the microbiome contributes to a specific disease or health in general.

Integrating data across investigations through a meta-analysis is a proven tactic for overcoming these limitations[13–15]. Though their application in microbiome science remains limited, meta-analyses provide important clarity in microbiome research. For example, meta-analysis of 16S-based investigations surrounding human obesity revealed that originally reported associations between the taxonomic composition of the gut microbiome and obesity were inconsistent across studies[16] and appear to only manifest weak statistical effects[17]. Additionally, a meta-analysis of 16S data quantified the microbiome’s taxonomic association with disease across several populations that span a variety of diseases revealing that some microbiome changes are disease specific while others are associated with multiple diseases[14]. The application of meta-analyses to shotgun metagenomic data is even more restricted, in part due to the limited amount of clinical metagenomic data currently available. One study integrated metagenomes to assess the predictive capacity of the taxonomic profile of the microbiome for several diseases finding that integrating multiple datasets improved prediction capabilities[15]. These studies highlight the importance of data integration in contributing to our understanding of the role of the microbiome in health and disease.

While these studies have proven insightful, their focus on taxonomy may limit our understanding of how the microbiome relates to health. Metagenomes afford insight into the types of genes, and consequent biological pathways, encoded by the microbiome. Resolving the association between microbiome functions and health may prove critical to determining the mechanisms through which the microbiome promotes health or contributes to diseases and may reveal robust indicators of disease given observations of functional redundancy where different microbes can fulfill the same functional roles[18, 19]. However, the application of meta-analyses to the functional diversity of the gut microbiome is limited to a single study of Type 2 diabetes which revealed gene families encoded in the microbiome that consistently associate with diseases across two continents[20]. The integration of metagenomic datasets in this study revealed the confounding contribution of anti-diabetic medicine to the results, emphasizing the need to consider additional factors, such as medication, in assessments of the gut microbiome’s relationship to health and disease.

Here, we describe the first meta-analysis of microbiome gene functions that span multiple disease-types and populations. We integrated ∼2,000 publically available metagenomes that span 8 studies and 7 diseases through a regression-based statistical framework. Our study (1) reveals that functional diversity indicates disease, but usually with weak effect, (2) resolves microbiome functions that associate with multiple diseases, suggesting that these functions may be important to host health in general, (3) identifies functions that indicate specific diseases, which could aid the development of diagnostics, and (4) quantifies the large effect of study-specific parameters on the functional diversity of the gut microbiome.

## METHODS

### Dataset

Our analysis relied on public metagenomes, which we obtained from the NCBI SRA and identified using SRAdb[21]. Specifically, we downloaded 10 Tbp of metagenomic sequence data from 1,979 subjects across 8 studies and 5 countries. Included in the dataset are non-diseased controls as well as subjects with one or more of the following diseases: rheumatoid arthritis[10], colorectal cancer[5], liver cirrhosis[8], Crohn’s disease [9, 11], obesity[5–12], type II diabetes[5–7,11,12], and ulcerative colitis[9, 11] (Table 1). Sample covariates were obtained from the initial studies.

**Table1:** Overview of case and control populations. The summary of case/control counts, subject age range, and source of the subjects summarized below for each of the seven diseases analyzed.

| Disease | Disease Level | Cases | Controls | Age Range | Studies |
| --- | --- | --- | --- | --- | --- |
| Rheumatoid Arthritis | Moderate | 45 | 93 | 19-70 | [10] |
|  | High | 70 |  |  |  |
| Colorectal Cancer | Advanced adenoma | 28 | 35 | 43-86 | [5] |
|  | Carcinoma | 36 |  |  |  |
| Liver Cirrhosis | - | 120 | 114 | 18-78 | [8] |
| Crohn's Disease | - | 12 | 132 | 18-70 | [9,11] |
| Obesity | - | 211 | 684 | 14-75 | [5,6,8–12] |
| Type II Diabetes | - | 204 | 366 | 13-86 | [5–7,11,12] |
| Ulcerative colitis | - | 58 | 132 | 18-70 | [9,11] |

### Data processing and annotation

We applied the standard MetaQuery quality control steps to quality filter all data and remove sequencing runs that failed quality control. Specifically, we deemed runs to be of sufficient quality if all of the following conditions were met: > 70% of reads mapped to the reference database, the mean read length was > 50-bp, the mean GC-content was > 40%, the mean read quality was > 21, and > 98% of reads were free of adaptor contamination. This left us with 3846/4230 (91%) of sequencing runs and 1885/1977 (95%) of subjects.

High-quality data was subject to functional annotation via MetaQuery[22] to produce profiles of genomic copy number of KEGG Orthology Groups (KOs) in each metagenome. Briefly, MetaQuery uses Bowtie2[23] to align sample reads to an integrated gene catalog for the human gut metagenome[7] to produce coverage estimates for each gene, which are functionally annotated by KOs. The coverage estimates are normalized by a set of 30 universal single copy genes[24] to produce estimates of the genomic copy number of each KO in the microbiome.

### Case and control population sampling

For each disease, we identified a set of subjects that represented disease-afflicted individuals (i.e. cases) and a set of subjects that represented healthy individuals (i.e. controls) (Table 1). Specifically, we deemed subjects that were clearly determined to have the disease of interest and none of the other diseases in the available metadata to be case subjects (i.e., no co-morbidity). Control populations alternatively consisted of individuals who were explicitly determined through clinical screening not to have the disease of interest, irrespective of the specific study from which the metagenome was generated. Designation of cases and controls relied on the metadata provided by the initial study. It is possible that a subject could have an undetected disease that was not screened for in a given a study. Rheumatoid arthritis subjects that manifested low disease severity or remission were excluded from these analyses. Subjects whose metagenomes were sequenced multiple times, either as technical or biological replicates, were only represented by the first metagenome sample that the study authors generated. Ultimately, 1,473 samples passed these analytical filters and were included in the downstream analyses.

### Alpha and beta-diversity

The vegan package in R was used for alpha and beta-diversity quantification. Specifically, the function specnumber (vegan::specnumber) assessed gene family richness and a two-sided t-test (stats::t.test) determined statistical significance between cases and controls within a disease. Beta-diversity was measured with Bray-Curtis dissimilarity (vegan::vegdist) and visualized with non-metric multidimensional scaling (NMDS). Permutational Multivariate Analysis of Variance (vegan::Adonis) calculated significant differences in beta-diversity. Beta-dispersion was quantified with betadisper (vegan::betadisper) and analysis of variance (ANOVA,stats::anova) determined significant differences.

### Identifying functions and taxa that stratify cases and control

A regression-based approach modeled KEGG module abundances across populations to identify the functions that stratify cases and controls for each disease. To reduce dimensionality of the data, KOs were collapsed into modules and only modules with prevalence greater than 0.5 were tested. The model (cplm::cpglm) implements a Tweedie compound Poisson distribution with a degenerate distribution at the origin and a continuous distribution on the positive real line to appropriately model data where there are zeros but the values are otherwise continuous [25, 26]. For each functional module, the genomic copy number was used as the response variable and disease status as the predictor. For disease phenotypes with data from multiple studies, the source study was also included as a covariate in the models to account for study effects. False discovery rate correction (stats::p.adjust) was used to adjust for multiple tests and a cutoff of FDR < 0.2 was used to identify indicators for each disease.

## RESULTS

### Gut metagenome functional diversity associates with disease

We integrated publically available gut metagenomic data from 1,473 patients spanning seven diseases and eight studies to discern how the functional diversity of the gut microbiome associates with disease. In particular, we investigated how protein family richness, the functional composition of the gut microbiome, and gut microbiome functional beta-dispersion relate to disease. These analyses clarify the diagnostic potential of gut microbiome functional diversity and point to putative microbiome-mediated disease mechanisms.

We observe a decrease in the number of KEGG Orthology Groups encoded in the gut metagenome (i.e., protein family richness) for four of the seven disease-afflicted populations relative to their respective control populations (Figure 1). Specifically, individuals diagnosed with Crohn’s disease (p < 0.001), obesity (p < 0.05), type II diabetes (p < 0.05), or ulcerative colitis (p < 0.01) manifest reduced gut microbiome protein family richness. In contrast, microbiome protein family richness is higher in subjects with colorectal cancer relative to their controls (p < 0.01). The protein family richness in the microbiome of subjects with liver cirrhosis or rheumatoid arthritis is similar to their respective controls. Our findings expand upon prior observations [11,27,28] and suggest that many diseases result in signatures of reduced protein family richness. However, the effect size of these differences in richness are generally miniscule, indicating that protein richness may not be an effective clinical diagnostic.

**Figure 1:**
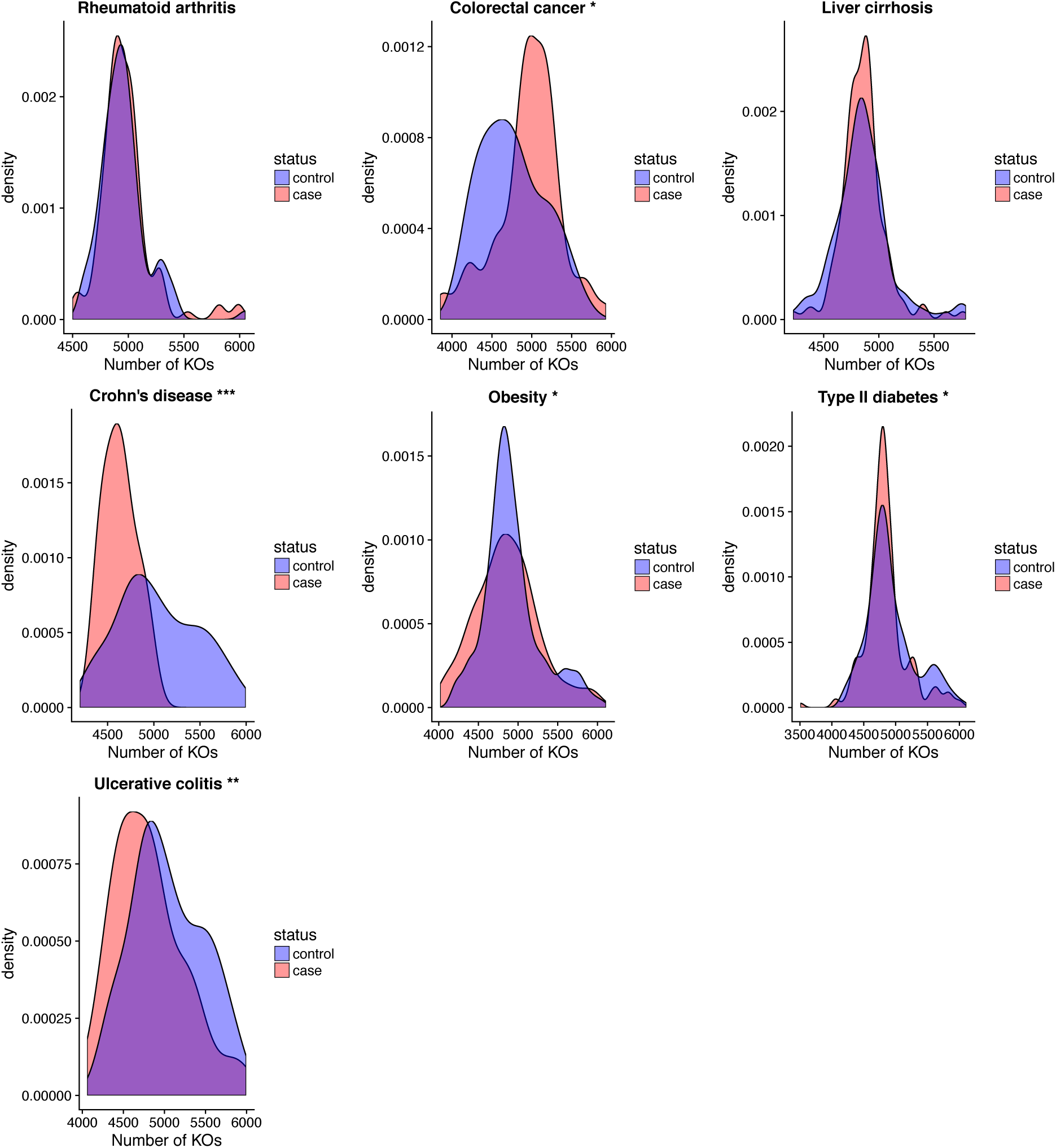
Protein family richness associates with disease. Density plots of the distribution of protein family richness across case and control populations for the seven diseases. Asterisks beside plot title indicates significance from Student’s t-Test (* p<0.05, ** p<0.01, *** p<0.001). Similar results were observed with Kolmogorov-Smirnov and Kruskal-Wallis tests.

Additional insight into how the gut microbiome relates to disease can be gleaned by quantifying the variation of the functional composition of the gut metagenome. We assessed this variation by measuring the beta-diversity of KEGG Orthology groups encoded in the gut metagenome using an abundance weighted metric (Bray-Curtis dissimilarity). Our analysis finds that the functional composition of the gut microbiome differs between case and control populations for the following five diseases: liver cirrhosis, Crohn’s disease, ulcerative colitis, obesity, and type II diabetes (Adonis p < 0.02; Figure 2). However, the magnitude of these differences varies across diseases, ranging from relatively strong effects in Crohn’s disease (partial R^2^ = 13%) to weak effects in obesity (partial R^2^ = 2%). Moreover, colorectal cancer and rheumatoid arthritis exhibit no detectable differences in functional beta-diversity between cases and controls. Similar to our richness analysis, these results indicate that the gut microbiome tends to encode different kinds of functions in association with disease, but that the differentiation is relatively modest and may challenge diagnosis.

**Figure 2:**
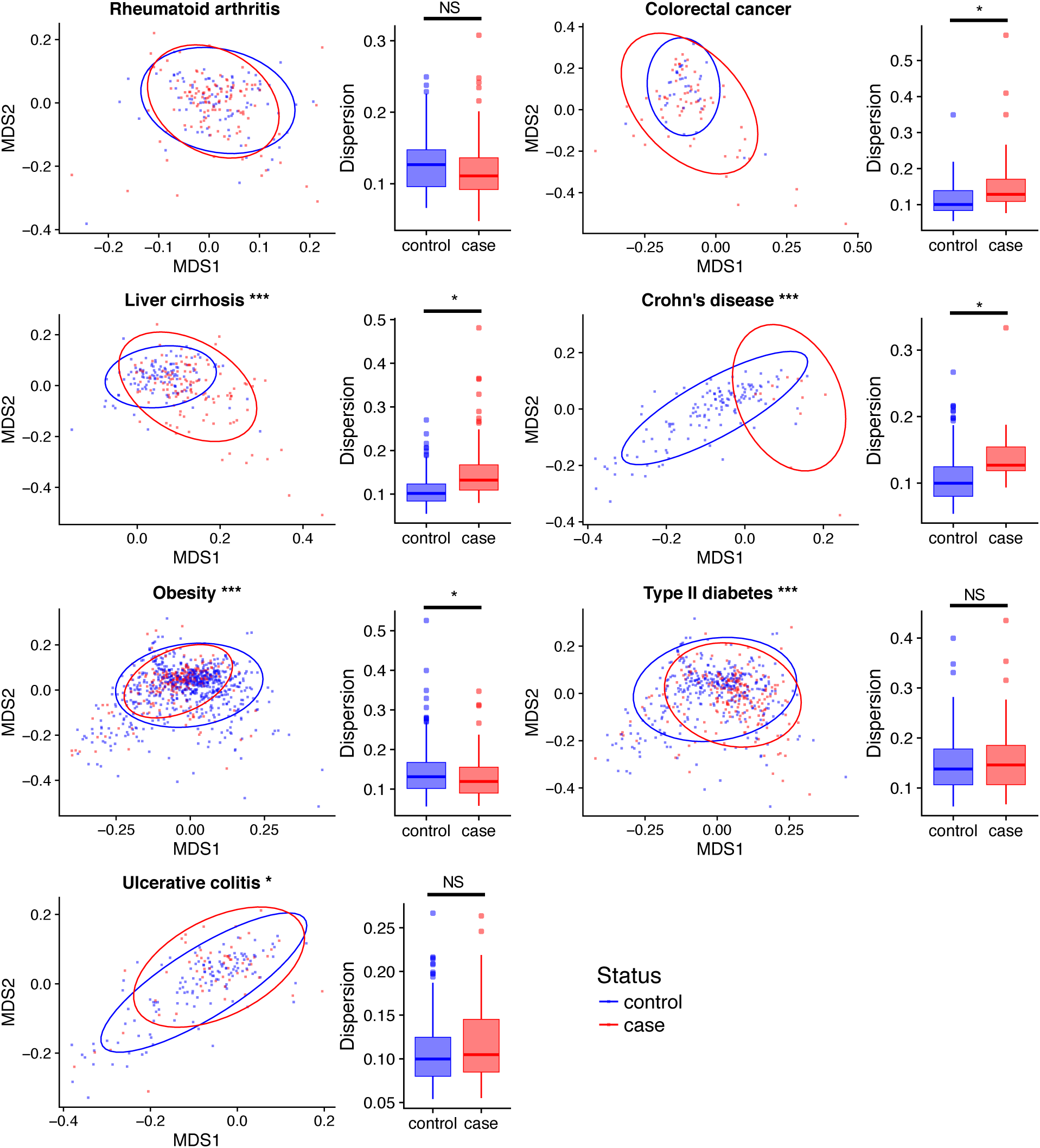
Changes in functional composition associates with disease. NMDS plots of Bray-Curtis dissimilarity between cases and controls across diseases. Asterisk in NMDS plot title indicates significance from PERMANOVA (*** p < 0.001,* p <0.05). Boxplots represent dispersion in beta-diversity within groups. Asterisk denotes significance from p-test and ANOVA (* p <0.05).

Beta-dispersion measures the compositional variation of the microbiome among a group of individuals, and prior work has linked disease to an increase in taxonomic beta-dispersion [29]. We similarly measured whether gut microbiome functional beta-dispersion varies between healthy and diseased populations. We observe an increase in functional beta-dispersion among individuals diagnosed with colorectal cancer, Crohn’s disease, and liver cirrhosis (p < 0.05; Figure 2). Individuals afflicted with obesity display a reduced beta-dispersion relative to their controls, but these results could be confounded by study effects. The remaining diseases presented no detectable difference in functional beta-dispersion. As observed with functional richness and beta-diversity, the effect size of beta-dispersion varies across diseases, but for some diseases appears to be relatively substantial. These results indicate that several diseases associate with an increased variation in the functional composition of the gut microbiome, possibly because there are relatively few functional assemblages that associate with control populations and many functional assemblages linked to disease.

### Metagenome modules indicate disease and clarify mechanisms of health

We next examined whether specific microbiome functions associate with disease. To reduce data dimensionality, we collapsed KEGG Orthology Groups into modules. We then used compound Poisson linear regression to model the relationship between health state and the average genomic copy number of KEGG modules. These methods have been applied in prior work [26] and allow for robust modeling of sparse but otherwise continuous data. Moreover, they afford the ability to account for potential study effects through the inclusion of additional covariates. Using these models, we defined indicators of a disease to be those modules whose average genomic copy number in the metagenome significantly associates with the health status of the host.

We find that of the 521 modules defined across our data set, 484 indicated disease in one or more of the disease populations (FDR < 0.2). The number of modules that indicate disease varied considerably across diseases. For example, 333 and 349 modules respectively indicate liver cirrhosis and Crohn’s disease, while only 13 modules indicate Ulcerative colitis. These results are qualitatively consistent at lower FDR thresholds.

The vast majority of the disease-indicating modules act as indicators for multiple diseases. Specifically, 78% of the indicator modules associate with two or more diseases and all diseases associate with relatively few unique indicators. For example, liver cirrhosis and Crohn’s disease share 92.5% and 88.8% of their indicators with another disease (Figure 3, Table 2). These results suggest that different diseases may manifest similar mechanisms of association with the gut microbiome (e.g., inflammation), or that microbiome modules may play varying roles in determining how the microbiome associates with different diseases.

**Figure 3:**
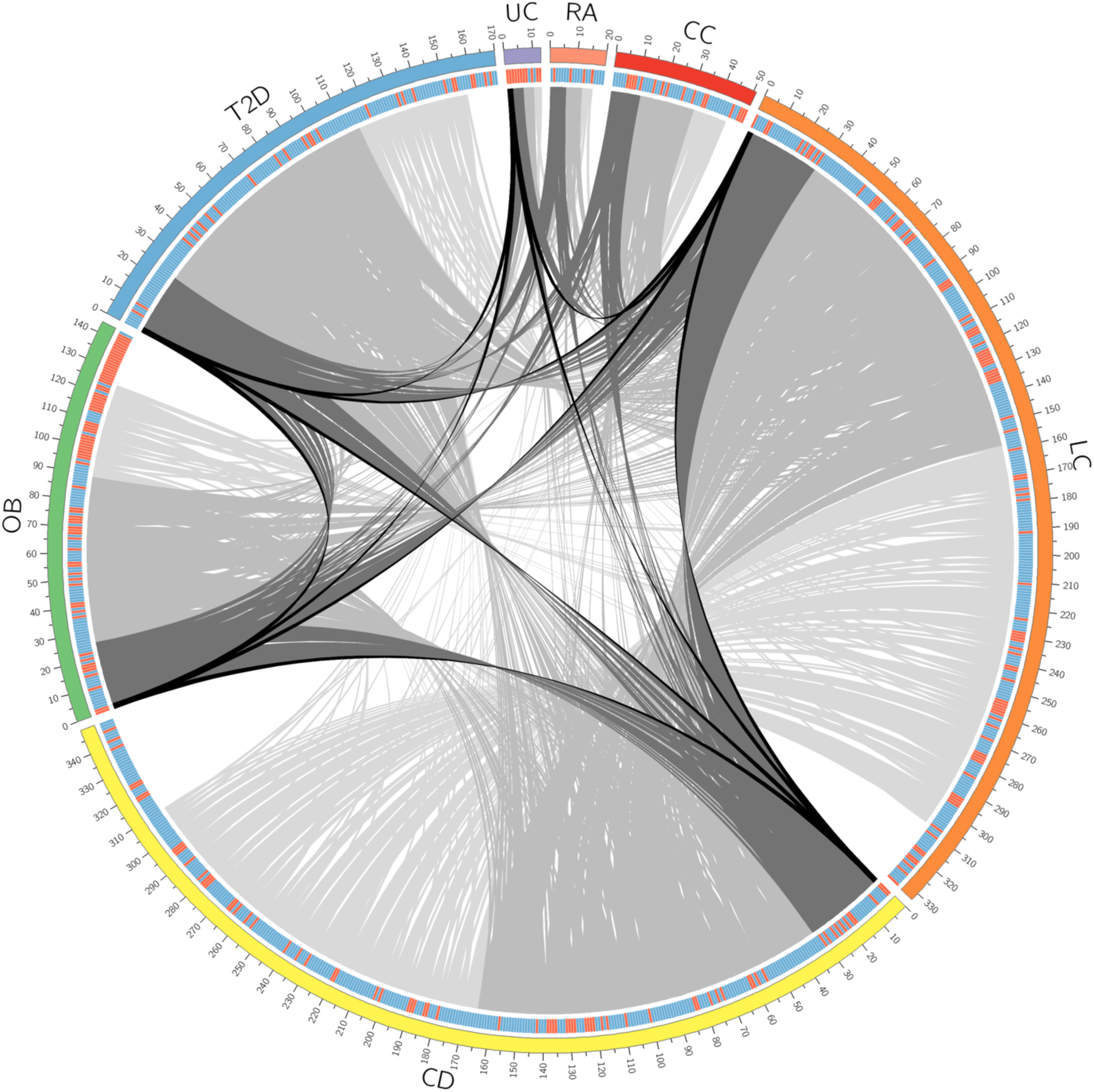
Most indicators of disease are shared between 2 or more diseases. Circos plot depicting overlap of indicator modules between diseases. Outer track represents the total number of module markers for each disease, tick marks represent counts of modules. Second track is a heat map with blue representing an increased abundance in cases and red representing a decrease in abundance in cases. Modules in each disease are ordered and links colored by the number of diseases they are indicators for (black = 5 diseases, dark grey = 4 diseases, grey = 3 diseases, light grey = 2 diseases). Modules without links are unique to the given disease. Abbreviations as follows: CC – colorectal cancer; LC – liver cirrhosis; CD, Crohn’s disease; OB – obesity, T2D – type II diabetes; UC – ulcerative colitis; RA – rheumatoid arthritis.

**Figure 4:**
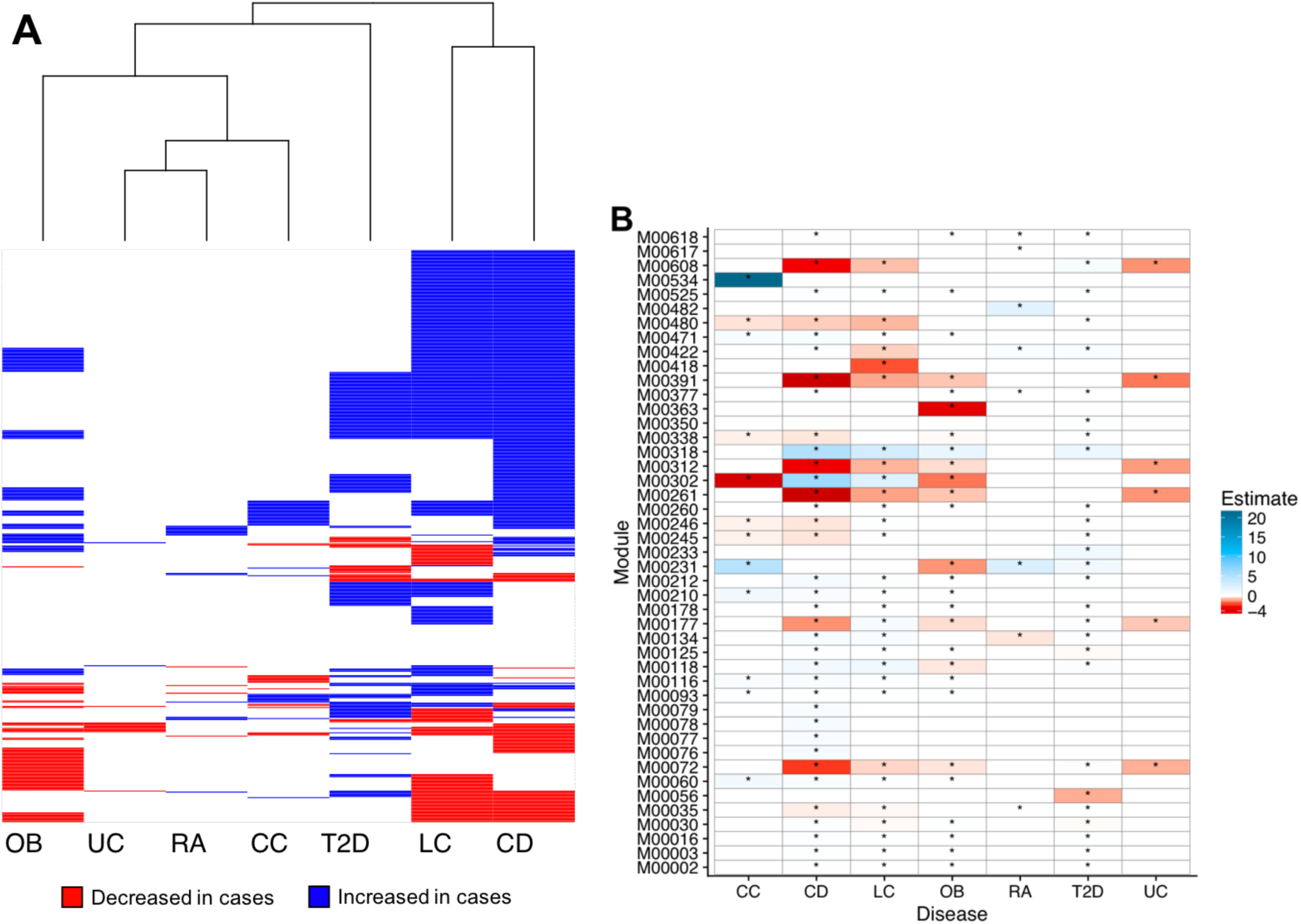
**A)** Heat map of KEGG modules colored by the sign of the slope estimate from the regression models. Blue indicates a positive slope estimate meaning higher abundance in cases and red indicates a negative slope estimate meaning lower abundance in cases (FDR < 0.2). White indicates no significant difference in module abundance for that disease. **B)** Heat map of highlighted common and unique indicators colored by the model slope estimate indicating the directionality and magnitude of the change in abundance. Asterisks denote significance (FDR < 0.2). Abbreviations as follows: CC – colorectal cancer; LC – liver cirrhosis; CD, Crohn’s disease; OB – obesity, T2D – type II diabetes; UC – ulcerative colitis; RA – rheumatoid arthritis.

**Table2:** Summary of module markers overlap between all pairs of diseases. Counts of modules that are differentially abundant between cases and controls for all pairs of diseases. Diagonal count represents the total marker count for each disease (FDR < 0.2)

|  | Crohn's Disease | Type II Diabetes | Colorectal Cancer | Liver Cirrhosis | Obesity | Ulcerative Colitis | Rheumatoid Arthritis |
| --- | --- | --- | --- | --- | --- | --- | --- |
| Crohn's Disease | **349** | 116 | 34 | 249 | 90 | 11 | 9 |
| Type II Diabetes | 116 | **170** | 15 | 119 | 38 | 3 | 8 |
| Colorectal Cancer | 34 | 15 | **51** | 26 | 11 | 0 | 2 |
| Liver Cirrhosis | 249 | 119 | 26 | **333** | 87 | 8 | 8 |
| Obesity | 90 | 38 | 11 | 87 | **144** | 9 | 7 |
| Ulcerative Colitis | 11 | 3 | 0 | 8 | 9 | **13** | 0 |
| Rheumatoid Arthritis | 9 | 8 | 2 | 8 | 7 | 0 | **20** |

We reasoned that the high frequency of modules that indicate multiple diseases may reflect the existence of modules that indicate any disease. While no modules stratified cases and controls across all seven diseases in our analysis, 33 modules indicated at least four but no more than five distinct diseases, and each disease is indicated by at least six of these 33 common disease indicators. Some diseases are indicated by a large proportion of these modules, including Crohn’s disease (97%), liver cirrhosis (88%), type II diabetes (73%), and obesity (79%). Conversely, rheumatoid arthritis (18%), colorectal cancer (33%), and ulcerative colitis (18%) are indicated by relatively few modules. That said, these modules can constitute a substantial fraction of the total indicators discovered for this latter set of diseases, as evidenced by the fact that 46% of the ulcerative colitis modules are among this common set. Despite the frequent number of diseases indicated by these modules, they do not always indicate diseases through consistent signatures. For example, N-glycosylation by oligosaccharyltransferase (M00072), which happens to indicate the largest number of diseases, is consistently depleted in individuals affected by liver cirrhosis, Crohn’s disease, obesity, type II diabetes, and ulcerative colitis relative to controls. Conversely, modules for cobalt/nickel transport systems (M00245 and M00246) are depleted in subjects with colorectal cancer and Crohn’s disease but elevated in subjects with liver cirrhosis and type II diabetes. Additional common indicators include modules associated with lipopolysaccharide biosynthesis and export (M00060, M00320, and M00080), iron/zinc/manganese/copper transport system (M00318), and acetate production (M00422, M00377, M00618, and M00579). These results demonstrate that while no microbiome modules universally signify health, there exist modules that are commonly perturbed during disease.

The relatively small number of modules that uniquely indicate disease provide insight into disease etiology and advance the development of disease specific diagnostics. Diseases varied in the proportion of their indicators that uniquely define the disease. For example, 20% of the rheumatoid arthritis indicators are unique while only 7% of type II diabetes indicators are unique. This observation highlights the fact that some diseases may offer greater potential for the discovery of microbiome-based clinical diagnostics. In rheumatoid arthritis subjects, the unique indicators include elevated levels of modules associated with methane production (M00617) and a DevS-DevR two-component regulatory system (M00482) that associates with *Mycobacterium tuberculosis* virulence [30]. The microbiomes of colorectal cancer subjects have increased abundance of a module for naphthalene degradation (M00534). The modules increased in the microbiome of liver cirrhosis subjects include nitrification (M00528) and staphylococcal virulence regulation (M00468). In contrast, there is a decrease in a module for toluene degradation (M00418). In Crohn’s disease, the modules increased in the microbiome of cases include degradation of glycosaminoglycans (M00076, M00077, M00078, and M00079) and B-vitamin biosynthesis (M00122, M00123, and M00573).

There is a decrease in abundance of modules associated with methanogenesis (M00576), antimicrobial peptide response (M00470), and phosphatidylethanolamine biosynthesis (M00092). The unique modules in the microbiome of obesity subjects include Enterohemorrhagic Escherichia coli (EHEC) pathogenicity signature (M00363). Type II diabetes cases have an increase in nitrogen fixation (M00175), glutamate transport (M00233), and capsaicin biosynthesis (M00350) and a decrease in O-glycan biosynthesis (M00056) in their gut microbiomes. There are not any unique indicators for subjects with ulcerative colitis.

### Study integration reveals robust functional indicators of obesity

Seven independent studies generated shotgun metagenomes for subjects with (n=211) and without (n=684) obesity. We integrated data across these studies by adding a study covariate to our regression models to resolve indicators of disease that are robust to study effects. We find that the source study accounts for approximately 25% of the variation in functional composition between obese individuals relative to controls, while disease status accounts for only 0.7%. Despite the contribution of study effects, we were able to detect 144 KEGG modules that indicate disease in the microbiome of subjects with obesity. This set of obesity indicators includes thiamin biosynthesis (M00127), EHEC pathogenicity signature (M00363), and acetate production (M00618, M00597, M00377).

## DISCUSSION

Our integrative analysis reveals the functional attributes of the gut metagenome that relate to human health and disease. We show that for several diseases, healthy microbiomes encode higher protein family richness, significantly different functional compositions, and increased constraint on the variation in that composition as compared to disease-associated microbiomes. However, effect sizes are frequently weak and not all diseases manifest these trends. Moreover, we identify specific functional modules that associate broadly with disease and, therefore, may be important to maintaining host health. Additionally, we resolve disease-specific markers that help clarify disease etiology and may serve as potential diagnostic biomarkers. Ultimately, the microbiome functions that we identify as being enriched in healthy individuals and disrupted in diseased individuals may illuminate how the microbiome contributes to host health.

We find that disease tends to associate with a reduction in the number of distinct protein families encoded in the microbiome. One potential explanation for this observation is that disease results in an altered gut environment that places selective pressures on microbes to possess specific functions. The microbes that carry these functions may also have similarities in the remainder of their genomes, thereby reducing the overall functional diversity in the microbiome. Additional support for this potential explanation comes in the form of our observation that diseases with arguably more severe intestinal symptoms (i.e. Crohn’s disease) display the largest deficit in richness while diseases with subtle intestinal symptoms (i.e. rheumatoid arthritis) have little difference in protein family richness between cases and controls. Our observation of decreased functional richness in diseased populations is consistent with reports that taxonomic diversity tends to be reduced in the microbiome of diseased populations relative to controls [11,27,28]. These reports prompted the disappearing microbiota hypothesis [31], which suggests that generational loss of ancestral members of the gut microbiota has adverse repercussions on host health. Based on our results, this hypothesis could be extended to include protein family richness, where loss of ancestral functions carried out by the gut microbiota may result in host disease.

Our integration of metagenomic data also enabled comparison of the differences in the gut microbiome’s functional composition across a variety of diseases. We find that while the microbiome’s functional composition associates with host health, the strength of the association substantially varies by disease and is generally relatively small. This suggests that disease is not defined by a substantial restructuring of the functional composition of the gut microbiome. Rather, if the microbiome contributes to diseases it does so through changes in the abundance of specific protein families, which may be different in each diseased subject. Consequently, health is not necessarily defined by the sum total of the functional capacity of the microbiome. Furthermore, analysis of the microbiome’s functional beta-dispersion reveals that most diseases have increased inter-sample variation in the microbiomes of the case populations relative to the microbiomes of the control populations. This pattern of increased dispersion in disease-associated microbiomes was previously observed in studies of taxonomic diversity and dubbed the Anna Karenina principle (AKP) [29]. AKP hypothesizes that certain stressors elicit stochastic effects on the taxonomic composition of the microbiome to yield increased variation in the stressed group relative to control group. Our beta-dispersion analysis shows that AKP effects are also found in the functional profiles of the gut microbiome in diseased hosts. This observation indicates that the increased dispersion observed in the taxonomic analysis of diseased microbiomes is unlikely to be the result of redundant functional compositions across communities, since if that was the case we would expect to find little to no increase in dispersion in the functional profiles. That said, our observation does not preclude the possibility that different taxa encode a small set of redundant proteins that associate with the disease state. Additionally, our finding that there tends to be lower functional dispersion among healthy individuals indicates that there may exist greater constraints on how the microbiome operates among healthy individuals.

Our robust and integrative modeling approach reveals specific associations between microbiome function and health by identifying commonly perturbed functions that may be impactful to host health. Interestingly, most of the common indicators (i.e., indicators of four or more diseases) are increased in abundance in the microbiomes of diseased subjects relative to the microbiomes of control subjects. This suggests that the shared disease associations with the microbiome may be due to the elevated presence of some microbiome functions rather than their loss in the microbiome. For example, subjects with colorectal cancer, liver cirrhosis, Crohn’s disease, and obesity have increased abundance of a module for lipopolysaccharide (LPS) biosynthesis (M00060). LPS is a well-known pro-inflammatory molecule; increased LPS biosynthesis by gut microbiota could contribute to intestinal inflammation observed in subjects with these diseases. Additionally, some common indicators may clarify collective features of the intestinal environment across disease. For example, several modules for iron transport (M00318, M00190, M00240, M00243, M00317, M00319) are increased in the microbiomes of subjects with Crohn’s disease, liver cirrhosis, obesity, and type II diabetes. Iron is an important cofactor for both humans and microbes and is often the subject of conflict between host and pathogen [32]. Subjects with Crohn’s disease often have low iron levels due to reduced iron absorption and iron loss through intestinal bleeding [33]. Patients with obesity and liver cirrhosis also tend to have a higher risk for low iron levels [34, 35], while subjects with type II diabetes tend to have high iron levels [36]. Differing abundance of iron transporters in the gut microbiome is a consistent indicator for disease, but the relationship to etiology seems to differ by disease. Another common indicator is acetate production (M00377 and M00618), which is increased in the gut microbiome of subjects with rheumatoid arthritis, Crohn’s disease, obesity, and type II diabetes. Production of short chain fatty acids (SCFAs), particularly acetate and butyrate, are thought to act as signaling molecules between the gut microbiome and host and may play a role in host metabolism [37]. The finding that increased abundance of modules for acetate production is consistent across diseases supports prior work linking microbe-produced SCFAs to disease. Through integration of data spanning various host health states, we gain a clearer picture of how the microbiome operates in relation to host health and what constitutes a healthy microbiome. It is possible that there are reasons we don’t detect differences in these common modules for more diseases, such as insufficient sample size or inability to control for unknown sources of variation that mask meaningful differences. The lack of universal markers could be due to fundamental differences in the diseases being examined. Additionally, the indicators identified require further investigation to determine the association with disease as some indicators may be a response to disease while others may play a role in disease development or progression.

Another benefit to the integration of data from distinct diseases is the ability to disentangle disease-specific from common indicators, which can be used to clarify the etiology of specific diseases and potentially act as biomarkers for disease diagnosis. For example, rheumatoid arthritis cases have increased abundance of a module for methane production (M00618) relative to controls. Increased abundance of methane producing microorganisms and elevated levels of methane were reported in patients with multiple sclerosis, an autoimmune disease that affects the central nervous system [38]. The identification of this module as a unique indicator for rheumatoid arthritis here and similar findings in a related disease suggests that methane production by gut microbiota may associate with autoimmune conditions. Additionally, modules for degradation of glycosaminoglycans (GAGs) (M00076, M00077, M00078, M00079) are uniquely elevated in subjects with Crohn’s disease. Disruption of GAGs in Crohn’s disease subjects has been reported previously [39] and may be due to increased degradation by gut microbiota. The observation that this increased abundance of GAG degradation modules is unique to Crohn’s disease subjects suggests this could serve as a diagnostic indicator and may be a target for therapeutics to restore intestinal integrity.

While these disease-specific indicators suggest that the microbiome could be useful in diagnostics, a more accurate and reliable approach may be gained by moving away from categorical assessments of health and towards quantitative measures of health. Consideration of disease progression as well as additional physiological covariates may also improve the diagnostic potential of the gut microbiome. For example, these personalized properties of disease may explain the relatively high variation in microbiome composition amongst diseased individuals. Moreover, our results demonstrate that the development of microbiome-based diagnostics of disease must consider the sensitivity of the microbiome to study effects. Indeed, for the two diseases for which we could quantify study effects, we found that these effects are much larger than the effect of disease on the microbiome. This study-associated variance could come from a variety of sources, including subject recruitment, sample collection and processing, and sequencing technology. Regardless, future efforts to develop microbiome-based diagnostics should similarly strive to integrate data across diverse studies to resolve robust indicators of disease.

The observed indicators of disease also clarify the potential role of the microbiome in various diseases. By focusing on what the microbiome is capable of doing, rather than which taxa are present, and how this functional capacity associates with health, we can develop testable hypotheses about how the microbiome may mediate health and disease. For example, our work reveals robust associations between the functional composition of the gut microbiome and obesity. Among the indicators for obesity are modules for acetate production (M00377, M00579, M00618). Recent research connects acetate production by gut microbiota to metabolic syndrome via interaction with the host parasympathetic nervous system to promote insulin secretion [40]. These results are especially valuable in light of recent work that demonstrates an effect of the microbiome in obesity [41] but inconsistent [16] or weak [17] associations between the taxonomic composition of the gut microbiome and obesity. Notably, the overall functional diversity of the microbiome similarly manifests weak associations with obesity, but the aforementioned protein families robustly resolve the disease. Consequently, these specific indicators may serve as important leads in future studies of how the gut microbiome contributes to obesity and metabolic syndromes.

Collectively, our analysis discerns how the gut microbiome’s functional capacity relates to host health. Through integration of data spanning multiple health states, we observe broad patterns of microbiome changes in disease that clarify how the gut microbiome contributes to health. For example, the metabolic modules that are commonly perturbed during disease may reflect mechanisms through which the gut microbiome interacts with physiology to promote health. Future studies should explicitly test whether these microbiome functions are critical to maintaining health. Moreover, disease associates with a personalized alteration in the functional composition in the microbiome, as indicated by our beta-diversity and -dispersion analyses. This result indicates that microbiome-based therapies may need to consider patient-specific parameters to ensure efficacy. Additionally, we uncover disease-specific indicators that not only serve as diagnostic leads, but also clarify potential microbiome-mediated etiologies of disease. Future studies should similarly seek to test the effects of these microbiome functions on health. Ultimately, integrative data analysis can expand our understanding of the role of the microbiome in maintaining health, but requires more comprehensive patient data, standardized methodologies, and extended patient populations to maximize its utility.

## ACKNOWLEDGMENTS

We thank Jesse Zaneveld for insightful discussions on beta-dispersion, Svetlana Lyalina for guidance with compound-Poisson modeling, Andrey Morgun and Natalia Shulzhenko for helpful feedback.

## REFERENCES

1. Vuong HE, Yano JM, Fung TC, Hsiao EY. The Microbiome and Host Behavior. Annu Rev Neurosci [Internet]. Annual Reviews; 2017 [cited 2018 Mar 19];40:21–49. Available from: http://www.annualreviews.org/doi/10.1146/annurev-neuro-072116-031347

2. Knight R, Callewaert C, Marotz C, Hyde ER, Debelius JW, McDonald D, et al. The Microbiome and Human Biology. Annu Rev Genomics Hum Genet [Internet]. Annual Reviews; 2017 [cited 2018 Mar 21];18:65–86. Available from: http://www.annualreviews.org/doi/10.1146/annurev-genom-083115-022438

3. Cho I, Blaser MJ. The human microbiome: at the interface of health and disease. Nat Rev Genet [Internet]. Nature Publishing Group, a division of Macmillan Publishers Limited. All Rights Reserved.; 2012;13:260–70. Available from: http://dx.doi.org/10.1038/nrg3182

4. Huttenhower C, Gevers D, Knight R, Abubucker S, Badger JH, Chinwalla AT, et al. Structure, function and diversity of the healthy human microbiome. Nature [Internet]. Nature Publishing Group, a division of Macmillan Publishers Limited. All Rights Reserved.; 2012 [cited 2016 Nov 29];486:207–14. Available from: http://www.nature.com/doifinder/10.1038/nature11234

5. Feng Q, Liang S, Jia H, Stadlmayr A, Tang L, Lan Z, et al. Gut microbiome development along the colorectal adenoma-carcinoma sequence. Nat Commun [Internet]. Nature Publishing Group; 2015 [cited 2017 Apr 18];6:6528. Available from: http://www.nature.com/doifinder/10.1038/ncomms7528

6. Karlsson FH, Tremaroli V, Nookaew I, Bergstrom G, Behre CJ, Fagerberg B, et al. Gut metagenome in European women with normal, impaired and diabetic glucose control. Nature [Internet]. Nature Publishing Group, a division of Macmillan Publishers Limited. All Rights Reserved.; 2013 [cited 2017 Apr 18];498:99–103. Available from: http://www.nature.com/nature/journal/v498/n7452/full/nature12198.html?WT.ec_id=NATURE-20130606

7. Li J, Jia H, Cai X, Zhong H, Feng Q, Sunagawa S, et al. An integrated catalog of reference genes in the human gut microbiome. Nat Biotechnol [Internet]. Nature Publishing Group, a division of Macmillan Publishers Limited. All Rights Reserved.; 2014 [cited 2017 Apr 18];advance on:834–41. Available from: http://dx.doi.org/10.1038/nbt.2942

8. Qin N, Yang F, Li A, Prifti E, Chen YY, Shao L, et al. Alterations of the human gut microbiome in liver cirrhosis. Nature [Internet]. Nature Publishing Group; 2014 [cited 2017 Jan 4];513:59–64. Available from: http://www.nature.com/nature/journal/v513/n7516/full/nature13568.html#supplementary-information

9. Nielsen HB, Almeida M, Juncker AS, Rasmussen S, Li J, Sunagawa S, et al. Identification and assembly of genomes and genetic elements in complex metagenomic samples without using reference genomes. Nat Biotechnol [Internet]. Nature Publishing Group, a division of Macmillan Publishers Limited. All Rights Reserved.; 2014 [cited 2017 Apr 18];32:822–8. Available from: https://www.nature.com/nbt/journal/v32/n8/pdf/nbt.2939.pdf

10. Zhang X, Zhang D, Jia H, Feng Q, Wang D, Liang D, et al. The oral and gut microbiomes are perturbed in rheumatoid arthritis and partly normalized after treatment. Nat Med [Internet]. 2015;21:895–905. Available from: http://www.ncbi.nlm.nih.gov/pubmed/26214836 http://www.nature.com/nm/journal/v21/n8/full/nm.3914.html?WT.ec_id=NM-201508&spMailingID=49267143&spUserID=ODM2MjM4OTIwNAS2&spJobID=741033692&spReportId=NzQxMDMzNjkyS0

11. Le Chatelier E, Nielsen T, Qin J, Prifti E, Hildebrand F, Falony G, et al. Richness of human gut microbiome correlates with metabolic markers. Nature [Internet]. Nature Publishing Group; 2013 [cited 2017 Apr 18];500:541–6. Available from: http://www.nature.com/nature/journal/v500/n7464/full/nature12506.html?WT.ec_id=NATURE-20130829

12. Wang JJ, Qin J, Li Y, Cai Z, Li SS, Zhu J, et al. A metagenome-wide association study of gut microbiota in type 2 diabetes. Nature [Internet]. Nature Publishing Group, a division of Macmillan Publishers Limited. All Rights Reserved.; 2012 [cited 2017 Apr 18];490:55–60. Available from: http://dx.doi.org/10.1038/nature11450

13. Haidich AB. Meta-analysis in medical research. Hippokratia [Internet]. Hippokratio General Hospital of Thessaloniki; 2010 [cited 2018 Feb 19];14:29–37. Available from: http://www.ncbi.nlm.nih.gov/pubmed/21487488

14. Duvallet C, Gibbons SM, Gurry T, Irizarry RA, Alm EJ. Meta-analysis of gut microbiome studies identifies disease-specific and shared responses. Nat Commun [Internet]. 2017 [cited 2017 Dec 7];8. Available from: https://www.nature.com/articles/s41467-017-01973-8.pdf

15. Pasolli E, Truong DT, Malik F, Waldron L, Segata N, Cho I, et al. Machine Learning Meta-analysis of Large Metagenomic Datasets: Tools and Biological Insights. Eisen JA, editor. PLOS Comput Biol [Internet]. Public Library of Science; 2016 [cited 2017 Jan 10];12:e1004977. Available from: http://dx.plos.org/10.1371/journal.pcbi.1004977

16. Finucane MM, Sharpton TJ, Laurent TJ, Pollard KS. A taxonomic signature of obesity in the microbiome? Getting to the guts of the matter. Heimesaat MM, editor. PLoS One [Internet]. Public Library of Science; 2014 [cited 2017 Sep 15];9:e84689. Available from: http://dx.plos.org/10.1371/journal.pone.0084689

17. Sze MA, Schloss PD. Looking for a signal in the noise: Revisiting obesity and the microbiome. MBio [Internet]. 2016 [cited 2018 Feb 19];7. Available from: http://mbio.asm.org/content/7/4/e01018-16.full.pdf

18. Hubbell SP. Neutral theory and the evolution of ecological equivalence. Ecology [Internet]. 2006;87:1387–98. Available from: http://www.ncbi.nlm.nih.gov/pubmed/16869413

19. Turnbaugh PJ, Ley RE, Hamady M, Fraser-Liggett CM, Knight R, Gordon JI. The human microbiome project. Nature [Internet]. 2007 [cited 2018 Feb 28];449:804–10. Available from: http://www.pubmedcentral.nih.gov/articlerender.fcgi?artid=3709439&tool=pmcentrez&rendertype=abstract

20. Forslund K, Hildebrand F, Nielsen T, Falony G, Le Chatelier E, Sunagawa S, et al. Disentangling type 2 diabetes and metformin treatment signatures in the human gut microbiota. Nature [Internet]. 2015 [cited 2018 Feb 19];528:262–6. Available from: https://www.nature.com/articles/nature15766.pdf

21. Zhu Y, Stephens RM, Meltzer PS, Davis SR. SRAdb: query and use public next-generation sequencing data from within R. BMC Bioinformatics [Internet]. BioMed Central; 2013 [cited 2018 Mar 19];14:19. Available from: http://bmcbioinformatics.biomedcentral.com/articles/10.1186/1471-2105-14-19

22. Nayfach S, Fischbach MA, Pollard KS, K. A, S. A, S.F. A, et al. MetaQuery: a web server for rapid annotation and quantitative analysis of specific genes in the human gut microbiome. Bioinformatics [Internet]. Oxford University Press; 2015 [cited 2017 Feb 1];31:3368–70. Available from: https://academic.oup.com/bioinformatics/article-lookup/doi/10.1093/bioinformatics/btv382

23. Langmead B, Salzberg SL. Fast gapped-read alignment with Bowtie 2. Nat Methods [Internet]. NIH Public Access; 2012 [cited 2018 Mar 19];9:357–9. Available from: http://www.ncbi.nlm.nih.gov/pubmed/22388286

24. Nayfach S, Pollard KS. Average genome size estimation improves comparative metagenomics and sheds light on the functional ecology of the human microbiome. Genome Biol [Internet]. 2015 [cited 2017 Apr 26];16:51. Available from: http://genomebiology.com/2015/16/1/51

25. Zhang Y. Likelihood-based and Bayesian methods for Tweedie compound Poisson linear mixed models. Stat Comput [Internet]. Springer US; 2013 [cited 2017 Jan 16];23:743–57. Available from: http://link.springer.com/10.1007/s11222-012-9343-7

26. Sharpton T, Lyalina S, Luong J, Pham J, Deal EM, Armour C, et al. Development of Inflammatory Bowel Disease Is Linked to a Longitudinal Restructuring of the Gut Metagenome in Mice. Gilbert JA, editor. mSystems [Internet]. 2017 [cited 2017 Nov 18];2:e00036–17. Available from: http://msystems.asm.org/content/2/5/e00036-17.abstract

27. Erickson AR, Cantarel BL, Lamendella R, Darzi Y, Mongodin EF, Pan C, et al. Integrated metagenomics/metaproteomics reveals human host-microbiota signatures of Crohn’s disease. PLoS One [Internet]. Public Library of Science; 2012 [cited 2017 Nov 17];7:e49138. Available from: http://journals.plos.org/plosone/article?id=10.1371/journal.pone.0049138

28. Nayfach S, Bradley PH, Wyman SK, Laurent TJ, Williams A, Eisen JA, et al. Automated and Accurate Estimation of Gene Family Abundance from Shotgun Metagenomes. PLoS Comput Biol [Internet]. Cold Spring Harbor Labs Journals; 2015 [cited 2017 Apr 21];11:22335. Available from: http://journals.plos.org/ploscompbiol/article/file?id=10.1371/journal.pcbi.1004573&type=printable

29. Zaneveld JR, McMinds R, Thurber RV. Stress and stability: Applying the Anna Karenina principle to animal microbiomes [Internet]. Nat. Microbiol. 2017 [cited 2018 Jan 30]. Available from: http://www.nature.com.ezproxy.proxy.library.oregonstate.edu/articles/nmicrobiol2017121.pdf

30. Dasgupta N, Kapur V, Singh KK, Das TK, Sachdeva S, Jyothisri K, et al. Characterization of a two-component system, devR-devS, of Mycobacterium tuberculosis. Tuber Lung Dis [Internet]. 2000 [cited 2017 Jun 21];80:141–59. Available from: http://ac.els-cdn.com/S0962847900902405/1-s2.0-S0962847900902405-main.pdf?_tid=d3820e4a-56b2-11e7-b6e9-00000aacb362&acdnat=1498071351_925b81026fbf8924ce6643e64f5b8c81

31. Blaser MJ, Falkow S. What are the consequences of the disappearing human microbiota? Nat Rev Microbiol [Internet]. Nature Publishing Group; 2009 [cited 2018 Feb 28];7:887–94. Available from: http://www.ncbi.nlm.nih.gov/pubmed/19898491

32. Skaar EP. The battle for iron between bacterial pathogens and their vertebrate hosts. Madhani HD, editor. PLoS Pathog [Internet]. Public Library of Science; 2010 [cited 2018 Mar 2];6:1–2. Available from: http://dx.plos.org/10.1371/journal.ppat.1000949

33. Gasche C, Lomer MCE, Cavill I, Weiss G. Iron, anaemia, and inflammatory bowel diseases [Internet]. Gut. 2004 [cited 2018 Feb 28]. p. 1190–7. Available from: https://www.ncbi.nlm.nih.gov/pmc/articles/PMC1774131/pdf/gut05301190.pdf

34. Zhao L, Zhang X, Shen Y, Fang X, Wang Y, Wang F. Obesity and iron deficiency: A quantitative meta-analysis [Internet]. Obes. Rev. 2015 [cited 2018 Feb 28]. p. 1081–93. Available from: http://doi.wiley.com/10.1111/obr.12323

35. Gkamprela E, Deutsch M, Pectasides D. Iron deficiency anemia in chronic liver disease: Etiopathogenesis, diagnosis and treatment [Internet]. Ann. Gastroenterol. 2017 [cited 2018 Mar 2]. p. 405–13. Available from: www.annalsgastro.gr

36. Simcox JA, McClain DA. Iron and diabetes risk [Internet]. Cell Metab. 2013 [cited 2018 Mar 2]. p. 329–41. Available from: https://www.ncbi.nlm.nih.gov/pmc/articles/PMC3648340/pdf/nihms449221.pdf

37. Morrison DJ, Preston T. Formation of short chain fatty acids by the gut microbiota and their impact on human metabolism [Internet]. Gut Microbes Taylor & Francis; May 3, 2016 p. 189–200. Available from: http://www.ncbi.nlm.nih.gov/pubmed/26963409

38. Jangi S, Gandhi R, Cox LM, Li N, Von Glehn F, Yan R, et al. Alterations of the human gut microbiome in multiple sclerosis. Nat Commun [Internet]. Nature Publishing Group; 2016 [cited 2018 Jan 18];7:12015. Available from: http://www.nature.com/doifinder/10.1038/ncomms12015

39. Murch SH, MacDonald TT, Walker-Smith JA, Lionetti P, Levin M, Klein NJ. Disruption of sulphated glycosaminoglycans in intestinal inflammation. Lancet [Internet]. Elsevier; 1993 [cited 2018 Jan 18];341:711–4. Available from: http://www.sciencedirect.com/science/article/pii/014067369390485Y

40. Perry RJ, Peng L, Barry NA, Cline GW, Zhang D, Cardone RL, et al. Acetate mediates a microbiome–brain–β-cell axis to promote metabolic syndrome. Nature [Internet]. 2016 [cited 2017 Jun 23];534:213–7. Available from: https://www.ncbi.nlm.nih.gov/pmc/articles/PMC4922538/pdf/nihms786451.pdf

41. Turnbaugh PJ, Ley RE, Mahowald MA, Magrini V, Mardis ER, Gordon JI. An obesity-associated gut microbiome with increased capacity for energy harvest. Nature [Internet]. 2006 [cited 2017 Jun 8];444:1027–31. Available from: http://dx.doi.org/10.1038/nature05414

